# Chimpanzee and human ApoE isoforms differ in the stimulation of neurite differentiation consistent with structural predictions with relevance to brain development and aging

**DOI:** 10.1101/2025.05.21.655373

**Authors:** Max A. Thorwald, Mafalda Cacciottolo, Xiaogang Hou, Todd E. Morgan, Caleb E. Finch

## Abstract

**Background:** Among anthropoids, humans uniquely possess ApoE isoforms that modulate Alzheimer’s disease (AD) risk and brain aging. While chimpanzee and human ApoE4 share R112 and R158, chimps do not exhibit advanced AD. A key difference is T61 in chimps versus R61 in humans, structurally resembling ApoE3.

**Objective:** We examined how astrocyte-derived ApoE isoforms impact neuronal morphology and used structural modeling to explore functional divergence.

**Methods:** Neonatal rat hippocampal neurons were cultured with astrocyte-conditioned media (ACM) from mice expressing human ApoE3, ApoE4, or chimpanzee ApoE. Neuronal outgrowth was quantified after 72 hours.

**Results:** Chimpanzee ACM increased neurite number by 30% over human ApoE isoforms. However, chimpanzee ACM resembled ApoE4 functionally, producing 40% shorter neurites and spines. Structural modeling supported greater similarity to ApoE4.

**Conclusions:** Chimpanzee ApoE is structurally and functionally more similar to ApoE4 than ApoE3, revealing evolutionary distinctions relevant to AD risk and neurodevelopment.

## INTRODUCTION

The human ApoE alleles are unique among primates and most other mammals that are monogenic. Three decades ago, ApoE4 was hypothesized to have evolved from chimpanzee ApoE at least 6 million years ago. Chimpanzee ApoE differs from human by 9 amino acids; both share arginine R112 and R158.^1–5^ Chimpanzee ApoE from samples of wild-caught and captive have not shown coding variants, unlike in humans.^4^ ApoE3 is globally the most prevalent, with cysteine substituted for arginine at residue 112 of chimpanzee and spread in *H sapiens* within the last 200,000 years.^6^ Aging humans are also unique among the anthropoids in developing advanced neurodegeneration of Alzheimer’s disease (AD) and related disorders for which ApoE4 is a major genetic risk factor. Aging chimps acquire diffuse amyloid, but lack neuritic plaques with dystrophic synapses.^5,7,8^ This divergence may be due to an amino acid site that is critical for protein folding: human R61 versus chimpanzee T61.

Structural modeling and site-directed mutagenesis suggested that chimpanzee ApoE, although more similar to ApoE4 in sequence, may structurally resemble human ApoE3.^4^ The primary role of ApoE is for lipid transport and these few substitutions alter lipid binding affinity. ApoE3 is considered the more efficient lipid carrier with higher binding capacity than ApoE4; lipid binding capacity of chimpanzee ApoE iis not reported.^6,9^

In brain, astrocytes are the primary producers of ApoE. Astrocyte ApoE is lipidated predominantly with cholesterol and secreted for neuronal uptake particularly in the hippocampus, an AD relevant brain region.^10,11^ Astrocyte ApoE is involved in protecting neurons against fatty acid associated toxicity, supporting axonal growth, and synaptic density.^12–14^ ApoE impact on neuron maintenance is determined by the ApoE isoform where neurite outgrowth and spine formation are greater with ApoE3 than E4.^10,15,16^ The contribution of chimpanzee ApoE on neuron differentiation has not been studied and is relevant to brain development and age-related cognitive decline.^17–19^ Brain development also differs by ApoE allele in AD-relevant brain regions.^20^ The entorhinal cortex was thinner in ApoE4 carriers by gene dosage in healthy children and young adults aged 8-20 years.^21^ A subsequent study partially confirmed these large differences on cortical thickness.^22^

The current study explored the neurotrophic effects of astrocyte-derived ApoE3, ApoE4, and chimpanzee ApoE on the differentiation of primary rat neurons. Astrocyte-conditioned media (ACM) from immortalized astrocytes expressing each isoform was used to assess the ApoE isoform impact on primary neuron differentiation for neurite length, soma size, dendritic branching, and spine density (**Fig. 1A**). The astrocytes were cultured from mice with targeted replacement by chimpanzee ApoE.

**Figure 1:**
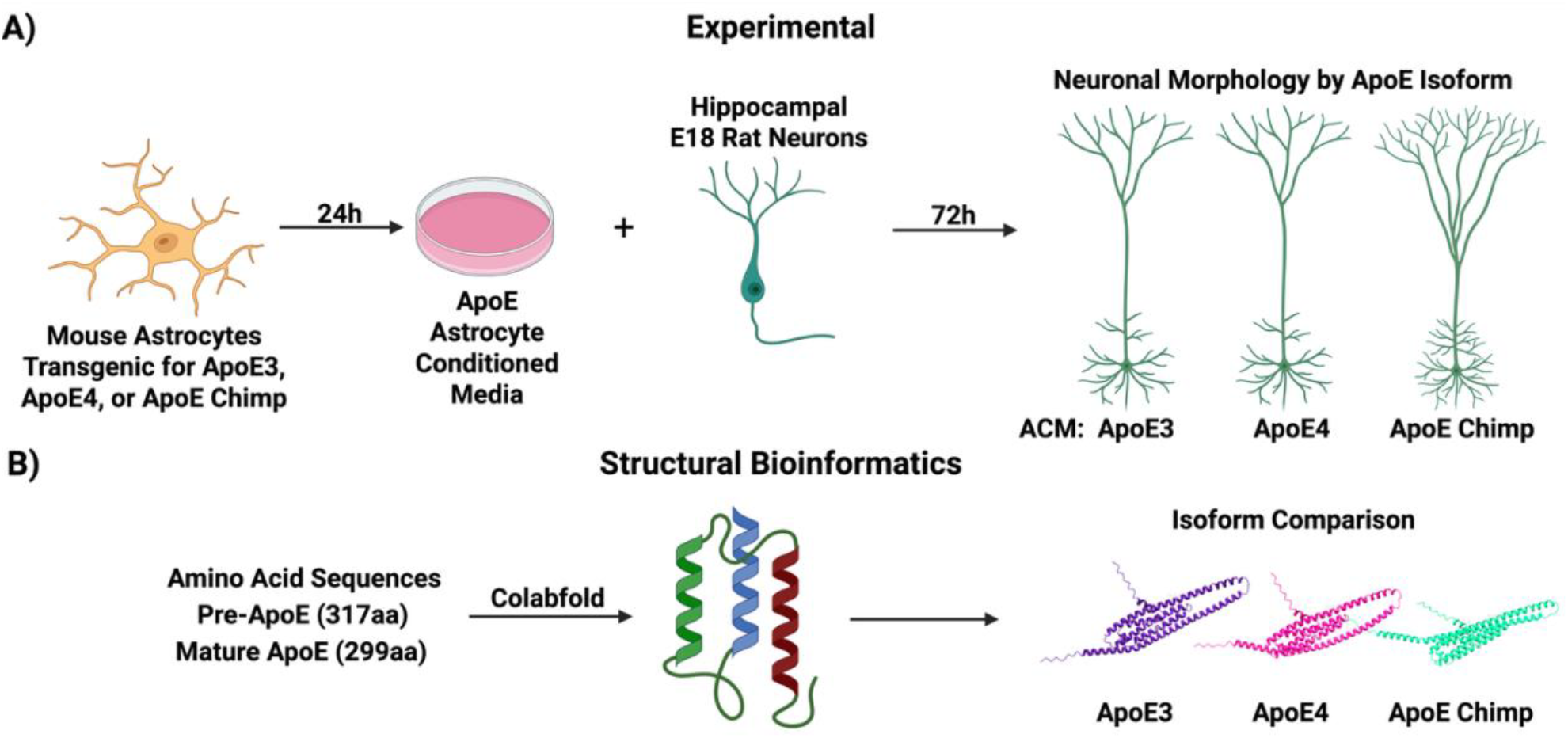
Study schematic. **A)** Astrocyte-conditioned media (ACM) was obtained from immortalized mixed glial culture from ApoE targeted-replacement mice. Day-18 embryonic hippocampal rat neurons were grown in ACM to study the effects of ApoE isoforms on neuron morphology. **B)** ApoE sequences were compared for structural differences by Colabfold.

We then generated structural folding predictions and extracted information of 3D space per residue of ApoE isoforms to determine if the amino acid substitutions altered critical regions such as those involved in lipid binding (**Fig. 1B**). We hypothesized that neurons exposed to chimpanzee ApoE in ACM would resemble those treated with ApoE3, consistent with its predicted structural similarity, but might diverge in function due to evolutionary adaptations. Neuronal characteristics that differed by ApoE isoform and species include neurite density and length, while structurally ApoE3 was more densely packed in lipid binding domains. We also confirmed that chimpanzee ApoE is more structurally similar to ApoE4, supported by *in vitro* data on neuron morphology. These findings identify further differences and similarities of chimp and human ApoE isoforms for their developmental roles; for the evolutionary context of AD vulnerability; and for the cellular underpinnings of neuronal morphology shaped by astrocytic lipid signaling.

## METHODS

### Chimpanzee ApoE targeted-replacement

Chimpanzee targeted-replacement mice (TR) were generated by replacing mouse ApoE with chimpanzee: **S.Fig. 1** compares human TR mice for ApoE3 and ApoE4.^15,16^ The chimpanzee ApoE gene, encompassing exons II-IV and associated introns, was subcloned from a chimpanzee bacterial artificial chromosome and inserted into a targeting vector, flanked by homologous mouse genomic sequences to facilitate homologous recombination. A neomycin resistance cassette provided positive selection; a diphtheria toxin A cassette was used for negative selection. The targeting vector was electroporated into embryonic stem (ES) cells from C57BL/6 mouse; transfected clones were selected using G418 (geneticin). Southern blot analysis screened for homologous recombination events. Genomic DNA was extracted from ES cell colonies and digested with EcoRI or HindIII, followed by hybridization with external and internal radioactively labeled probes. PCR-based screening confirmed the correctly targeted allele.

TR-ES cells were microinjected into blastocysts derived from C57BL/6 mice and implanted into pseudopregnant female mice. Chimeric offspring were identified by coat color mosaicism and then bred with C57BL/6 wild-type mice to obtain germline transmission of the targeted allele. Germline transmission was confirmed by PCR genotyping using primers specific for chimpanzee ApoE. RNA splicing analysis used RT-PCR to confirm appropriate expression and transcript integrity. After these studies were completed, the Chimp-TR line was lost; tissues from limited numbers of ages mice are available. Genomic DNA from the offspring was analyzed by Southern blotting to verify correct TR.

### ApoE Cell Lines

Immortalized primary astrocytes generated astrocyte condition media (ACM). Primary astrocytes from the neonatal ApoE chimp mice or ApoE3 and ApoE4 TR mice prior to immortalization.^24^ Cells were maintained in DMEM/F12 (Gibco #11330-032), 10% FBS, 1 mM sodium pyruvate, 200 µg/ml G418 and plated at 500k cells per dish. Upon confluency, cells were replated in serum-free media for 24 hours to obtain ACM. ApoE levels was measured by dot blot with an ApoE specific antibody (1:1000, goat, Millipore).

### Primary Neuronal Culture

Primary hippocampal neuronal cultures were derived from embryonic day 18 (E18) rats. Briefly, hippocampi were dissociated in Hank’s balanced salt medium containing trypsin and DNase at 37°C.^25^ Dissociated cells were plated on poly-D-lysine and laminin coated glass coverslips (20k cells/cm^2^), or on 96 well plates (70k cells/cm^2^). Cells were maintained in media DMEM supplemented with B27 (Invitrogen, Grand Island, NY), with serum-free media that favors neuronal survival.^3^ Neurons were maintained at 37°C with 5% CO_2_.

### Neuronal Morphology

E18 cultures of hippocampal neurons were treated with ACM for 72 hours. After incubation, cultures were fixed in 4% paraformaldehyde\phosphate buffered saline, pH 7.4 and immunostained for neuron-specific βIII-tubulin (1:500, rabbit; Sigma Chemical Co.) and F-actin in lamellipodia and growth cones (rhodamine phalloidin, 1:40; Molecular Probes). Images were captured by immunofluorescent microscopy using a Nikon Eclipse TE300 microscope (Nikon Inc., Melville, NY). One hundred or more neurons per ApoE isoform were sampled per measurement. Neuron morphology was analyzed in ImageJ. Neuronal length was assessed using the NeuronJ plugin in ImageJ.^27^

### ApoE Structural Analysis

Local folding of ApoE structures was done using Colabfold v1.5.5.^28^ ApoE3 and chimpanzee sequences were obtained from the Uniprot database. The ApoE3 sequence was modified to generate ApoE4. Folding predictions were done with model alphafold2_ptm with multiple sequence alignment for 12 recycles with amber relaxation. Five models were generated and ranked based on highest predicted local distance difference test (pLDDT) score; pLDDT scores per residue, **S.Table 1**. Forty-seven structures from NMR, x-ray crystallography, and cryo-EM were superimposed to assess prediction validity (**S.Table 2**). Secondary structures were assigned by DSSP (**S.Table 3**).^29^ Predicted structures were then subjected to Atom3D to extract nodes per residue represented as the amino acids within spatial proximity (≤10Å).^30^ Node features included atomic coordinates, amino acid identity, and backbone dihedral angles, while edge features encoded inter-residue distances and orientation vectors. Structures were visualized, aligned, and annotated with ChimeraX (ver.1.9) and color coded by data obtained from predicted structures or node connectivity.^31^

### Statistics

Group differences means (± SEM) were analyzed by one-way ANOVA with Tukey’s post-hoc test, Kruskal-Wallis with Dunn’s test for nonparametric distributions, or Welch’s ANOVA with Games-Howell test when variances were unequal across groups. Neuronal categorical outcomes (e.g., pyramidal vs. non-pyramidal or mature vs. immature neurons) were analyzed by Chi-square tests. Where applicable, significant contingency table results were followed by pairwise 2x2 comparisons with Bonferroni correction. Significance was defined as p < 0.05. Analyses used GraphPad Prism version 10 (GraphPad Software, San Diego, CA).

## RESULTS

### ApoE3 is associated with longer neurites and perikaryal area

To assess if ApoE-chimp isoform modifies neuron morphology differently than the two main human ApoE isoforms, embryonic rat day18 neurons were grown in astrocyte-conditioned media (ACM). Levels of the chimp and human ApoE isoforms did not differ in ACM by dotblot (**S.Fig. 2A**). Axonal length did not differ between neurons grown with ACM from ApoE4 or ApoE-chimp. However, ApoE3 ACM increased neuronal length by 35% above ApoE4 and ApoE-Chimp (**Fig. 2A**). ApoE4 and Chimp ACM treated neurons had fewer neurites longer than 200 μM than ApoE3 (**Fig. 2B**). The opposite was observed for the number of neurites per neuron, with ApoE-chimp having 25% more neurites (**Fig. 2C**). Perikaryal area was 40% greater in ApoE3 neurons than human E4 or Chimp (**Fig. 2D**).

**Figure 2:**
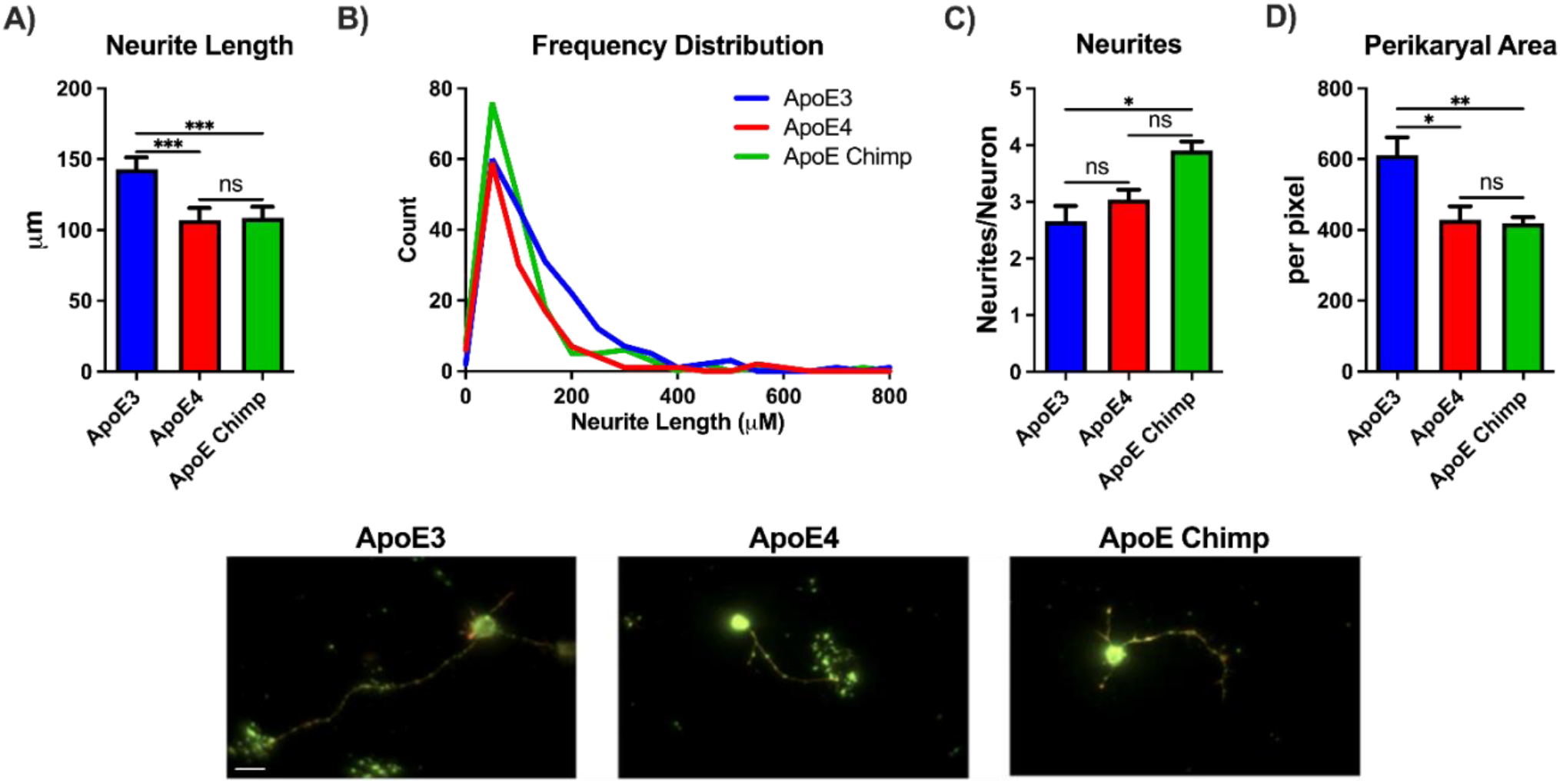
Neuronal morphology characterized by **A)** neurite length, **B)** frequency distribution of neurite length, **C)** neurites per neurons, and **D)** perikaryal area. Representative images of embryonic rat neurons grown 72 hours in ACM from ApoE3, ApoE4, or ApoE chimp ACM; scale bar, 15μM. Statistics by one-way ANOVA with Tukey’s posthoc or Kruskal-Wallis. *p<0.05, **p<0.01, ***p<0.001.

### ApoE3 is associated with greater neurite spine density and neuron maturity

Next, we examined the density of neuritic spines and levels of neuronal maturation. Neurons with ApoE3 ACM had 40% more spines than ApoE4 and 55% more than ApoE-Chimp. ApoE4 ACM treated neurons had 10% more spines than ApoE Chimp (**Fig. 3A,B**). Neuron maturity was evaluated by the type of spine present by classification: mature (mushroom or stubby) or immature (filopodia-like or thin).^32^ Neurons grown in ApoE3 ACM had 25% more mature forms than ApoE4. Chimp ApoE did not differ from ApoE3 and ApoE4 for neuron maturity (**Fig. 3C**).The type of neuron (pyramidal versus non-pyramidal) also did not differ by ApoE isoform (**S. Fig. 2B**).

**Figure 3:**
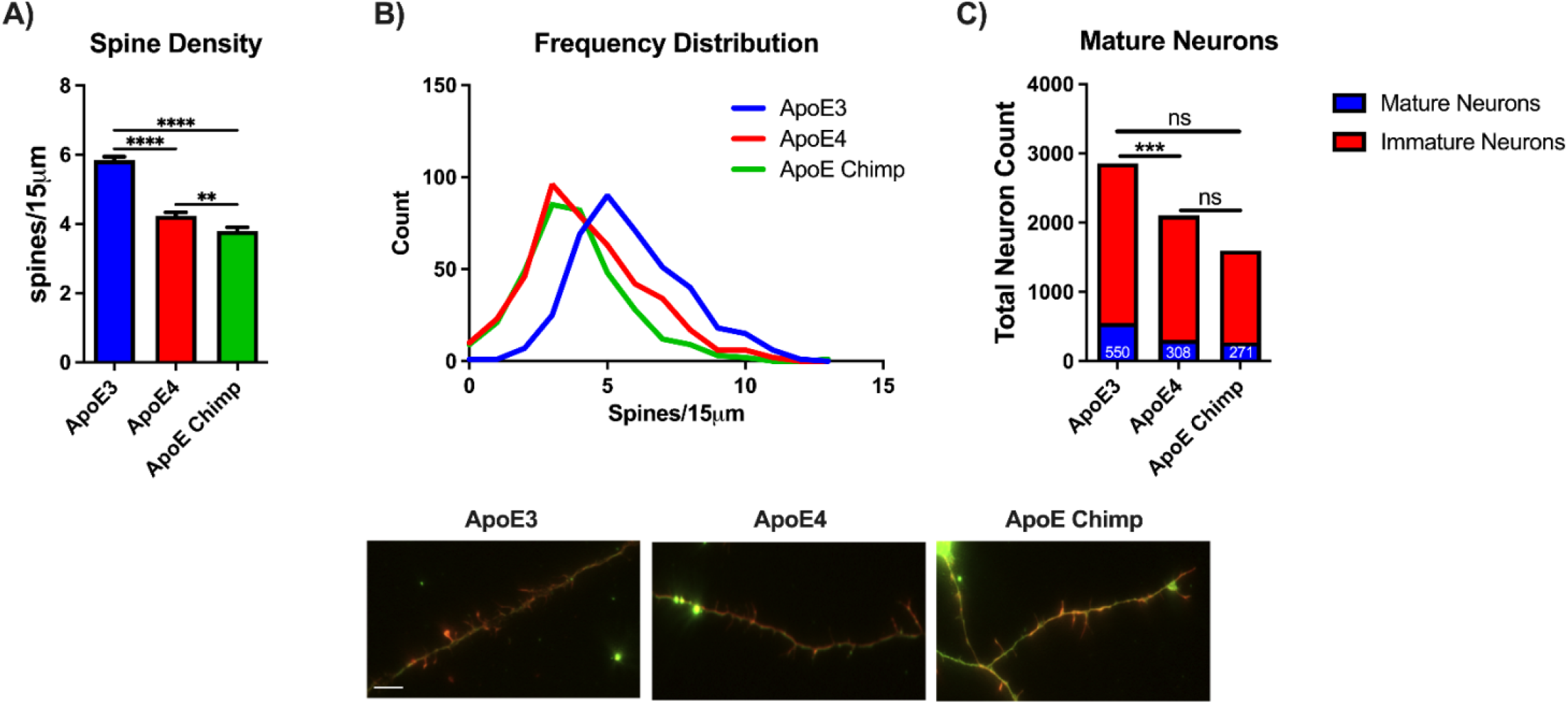
ApoE3 promotes the higher spine density. **A)** Neurite spine density, **B)** Frequency distribution of spine density, and **C)** Neuronal maturity of primary neurons grown in astrocyte-conditioned media (ACM) from ApoE-chimp, human ApoE3 and E4. Images of spine density in embryonic rat neurons exposed to ApoE3, ApoE4, or ApoE chimp ACM, 72 hours; scale bar 5μM. A) Statistics by one-way ANOVA with Tukey’s posthoc or Kruskal-Wallis. C) Statistics by Chi-square with Bonferroni. **p<0.01, ***p<0.001, ****p<0.0001. correction

### Chimpanzee ApoE is structurally more similar to ApoE4 than ApoE3

ApoE structure was modeled to quantify conformational differences associated with key isoform-defining residues (61, 112, and 158). Structures generated in Colabfold were compared to x-ray crystallography (**S. Fig. 3A**), nuclear magnetic resonance (NMR; **S.Table2**), and structures downloaded from AlphaFold’s database for the pruned and the full root mean standard deviation (RMSD) to measure structural similarity between proteins. Protein databases such as AlphaFold and UniProt used the ApoE precursor (pre-ApoE; 317aa). We found that the average pruned RMSD for our predicted structures and AlphaFold’s ApoE structure compared to the empirical structure data did not differ (**S.Fig. 3B**). However, comparison of the average full RMSD values with empirical data and the Colabfold precursor, mature, and AlphaFold showed our predicted pre-ApoE was 30% closer than our mature ApoE and 55% closer than AlphaFold’s model (**S. Fig 3C**).

After validating the predicted structures, we then analyzed pairwise structural superposition of the mature ApoE isoforms, which showed minimal geometric deviation between ApoE3 and ApoE4, with pruned RMSD of 0.273Å and full RMSD of 4.764Å. The pruned RMSD indicates strong conservation of the core fold, while the higher full RMSD suggests peripheral or loop region flexibility between isoforms. In contrast, alignment between ApoE3 and chimpanzee ApoE showed a larger pruned RMSD of 0.350Å and an even more pronounced full RMSD of 12.228Å, indicating greater global structural divergence (**Fig. 4**). The increased full RMSD reflects both localized atomic displacements and broader shifts in loop and domain positioning. Similarly, alignment between ApoE4 and chimpanzee ApoE revealed a pruned RMSD of 0.337Å and a full RMSD of 10.625Å (not shown).

**Figure 4:**
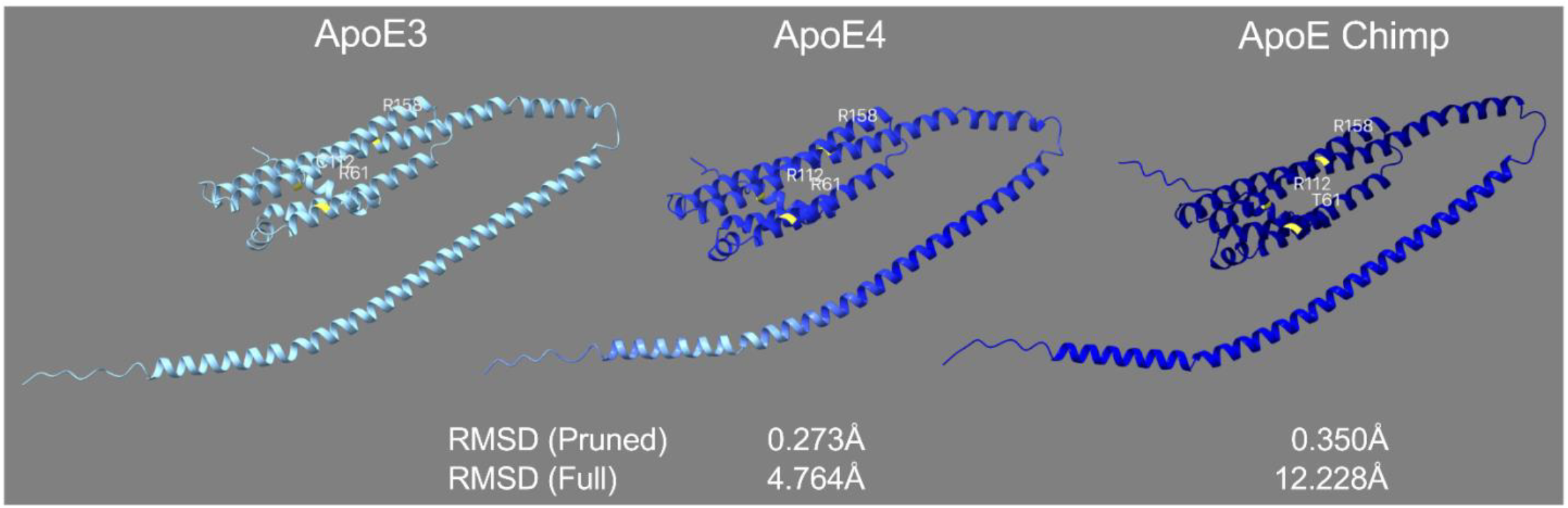
Comparison of mature ApoE3, ApoE4, and ApoE chimpanzee structures. Key residues at positions 61, 112, and 158 are highlighted in yellow. RMSD values (pruned and full) are calculated relative to ApoE3. Structures are colored by full RMSD gradient relative to ApoE3 to indicate structural deviation; darker indicates higher shifts in atom positions.

Pre-ApoE was then examined containing the 18 amino acid signal peptide. ApoE3 and ApoE4 had a pruned RMSD of 0.482Å and a full-length RMSD of 3.740Å. Pre-ApoE3 and chimpanzee ApoE again had larger pruned RMSD of 0.611Å and full RMSD of 6.611Å (**S.Fig 4**). ApoE4 and chimpanzee ApoE had a pruned RMSD of 0.625Å and a full RMSD of 5.506Å (not shown). Thus, while Pre-ApoE4 and chimpanzee ApoE are structurally more similar than ApoE3 in terms of overall folding, there are significant deviations, particularly in regions outside the core helices. In sum, the chimpanzee ApoE structure maintains a broadly similar core fold to human ApoE isoforms, but exhibits greater overall conformational divergence, particularly outside the core helix bundle.

### ApoE node connectivity is increased at position 61 for mature and precursor ApoE3

The further interrogate the influence of the few residue differences between ApoE isoform on structure we extracted per residue atom information from the predicted structures. The node connectivity (Δnode) is represented as the number of amino acids within a radius of 10Å from the reference amino acid. The total number of nodes per structure did not differ between the ApoE isoforms, mature or precursor, consistent with the few amino acid substitutions (**Fig. 5A, S.Fig.5A**). Nodes were plotted by amino acid to highlight regions that differed between the isoforms. For mature ApoE isoforms, position 112 had increased node connectivity (17 to 18) in ApoE3 compared to ApoE4 and Chimp. Position 158 had no node differences between isoforms, and positions 112 and 158 did not differ. Position 61 node connectivity was also increased in mature ApoE3 than ApoE4 and ApoE Chimp. For pre-ApoE both ApoE3 and ApoE4 had an increase in node connectivity with 18 atoms compared to 17 for chimp (**Fig. 5B, S.Fig. 5B**). These differences become more apparent when overlaying the Δnode degree on ApoE3 or ApoE4 backbones wherein blue indicates a loosening of the domain while red indicates a tighter structure. ApoE3 and ApoE4 are more similar for the mature forms with some differences in domain flexibility in the C-terminal side (**Fig. 5C**). Pre-ApoE3 and ApoE4 show moderate differences which become greater when comparing ApoE3 and ApoE-chimp. Conversely, comparing connectivity between pre-ApoE4 and ApoE Chimp, the connectivity differences are much smaller, consistent with the evolutionary trajectory (**S.Fig. 5C**). Interestingly, these differences are predominant in the lipid binding region (225-299; **Fig. 5B,C**). Thus, these few amino substitutions cause major conformational changes in domains essential to ApoE functions.

**Figure 5:**
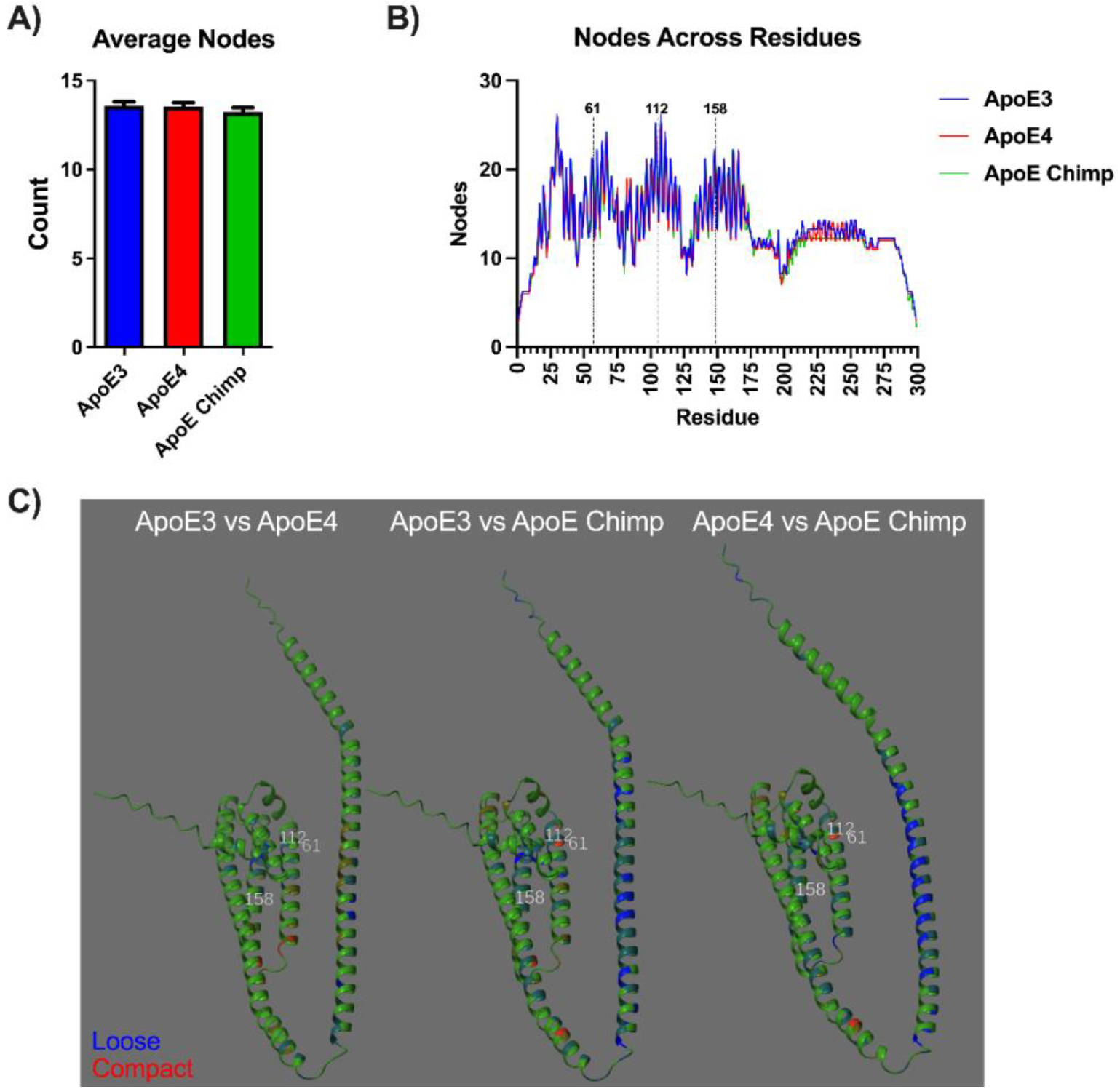
Amino acid connectivity per residue for precursor ApoE3, ApoE4, and chimpanzee ApoE. **A)** The average number of nodes per ApoE isoform within a 10Å radius. **B)** The number of nodes at each amino acid for precursor ApoE. Annotations (61, 112, and 158) are based on the mature 299 amino acid sequence. **C)** Reference structures of ApoE3 or ApoE4 are colored green by the Δnode degree for ApoE4 or ApoE chimpanzee. Red indicates increased node connectivity while blue represents decreased node connectivity.

**Table 1:**
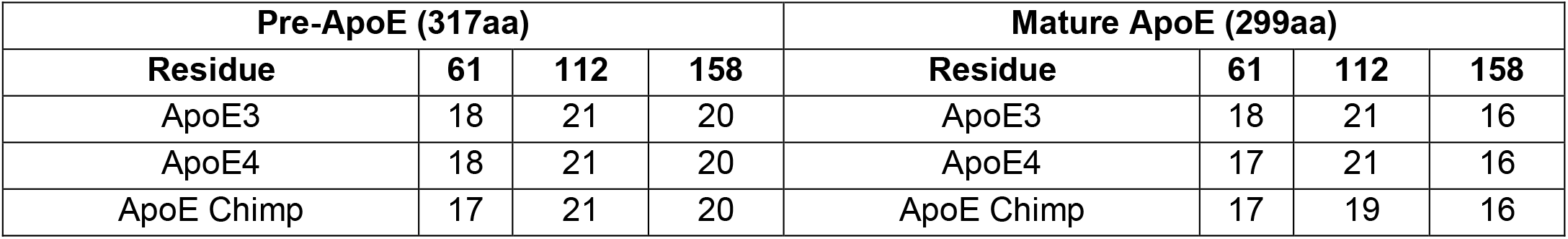
Node connectivity for pre-ApoE and mature ApoE at isoform defining residues.

## DISCUSSION

This first direct functional comparison of chimpanzee and human ApoE isoforms on neuronal development shows structural and morphological distinctions with implications for brain evolution and AD susceptibility. Using ACM derived from mice expressing human ApoE3, ApoE4, or chimpanzee ApoE, we assessed effects on primary hippocampal neuron morphology and linked it to high-resolution structural predictions. Contrary to former predictions that chimpanzee ApoE is functionally equivalent to ApoE3, we show that chimpanzee ApoE induces neuronal differentiation more similar to ApoE4, matching our structural predictions.^4^

Neurons exposed to chimpanzee ApoE ACM developed shorter neurites and reduced spine density than ApoE3, confirming prior studies on the restrictive influence of ApoE4 on neurite outgrowth and synaptic maturation.^33,34^ Unique to chimpanzee ApoE was the increased number of neurites per neuron, suggesting a divergent influence on early neurogenesis that may reflect adaptive evolutionary traits unrelated to AD vulnerability. Thus, while chimpanzee ApoE shares closer structural and sequence homology with ApoE4, its neurotrophic profile is not identical.

The structural analysis supports these functional differences. Chimpanzee ApoE shares greater overall similarity with ApoE4 than with ApoE3. Pruned RMSD, which reflects the core fold conservation, was modest across isoforms. Full RMSD values revealed substantial divergence between chimpanzee ApoE and ApoE3, suggesting domain-level flexibility and long-range conformational changes. Node-based residue connectivity analysis further confirmed that little difference exists between pre- and mature forms of ApoE for defining residues (112, and 158). However, 61 was associated with more connectivity with ApoE3 than ApoE chimp where ApoE matched ApoE3 for the mature structure. Overall, this may imply that functional divergence likely stems from global shifts in tertiary structure or domain reorganization rather than local residue environment alone. Furthermore, our data suggests that T61 alone does not recreate the neurotrophic properties of ApoE3 in the chimpanzee isoform. Instead, we observed broader structural loosening in chimpanzee ApoE than ApoE3, particularly at position 61, further decoupling local amino acid substitution from functional outcome. The pre-ApoE structural comparisons reinforced these findings, as chimpanzee ApoE maintained greater similarity to ApoE4 than to ApoE3 even before signal peptide cleavage, suggesting conserved folding trajectories during processing and secretion.

These phenotypic differences may be rooted in broader evolutionary divergence beyond the currently discussed residue positions (61, 112, 158). Noncoding regulatory elements, additional nonsynonymous substitutions within the ApoE gene, or adjacent loci on chromosome 19q13.32 which includes APOC1, APOC2, and APOC4 may contribute to the unique developmental and aging trajectories of the human brain (**Fig. 6**). In mice this cluster is found on 7qA1 consistent with the tracked translocation events from 90 million years ago.^35^ Recent studies suggest that these neighboring genes may modulate aspects of neuronal structure and AD risk independently or synergistically with ApoE.^36^ The present study did not consider contributions from other apolipoprotein family members within ACM which may also modulate neuron differentiation.

**Figure 6:**
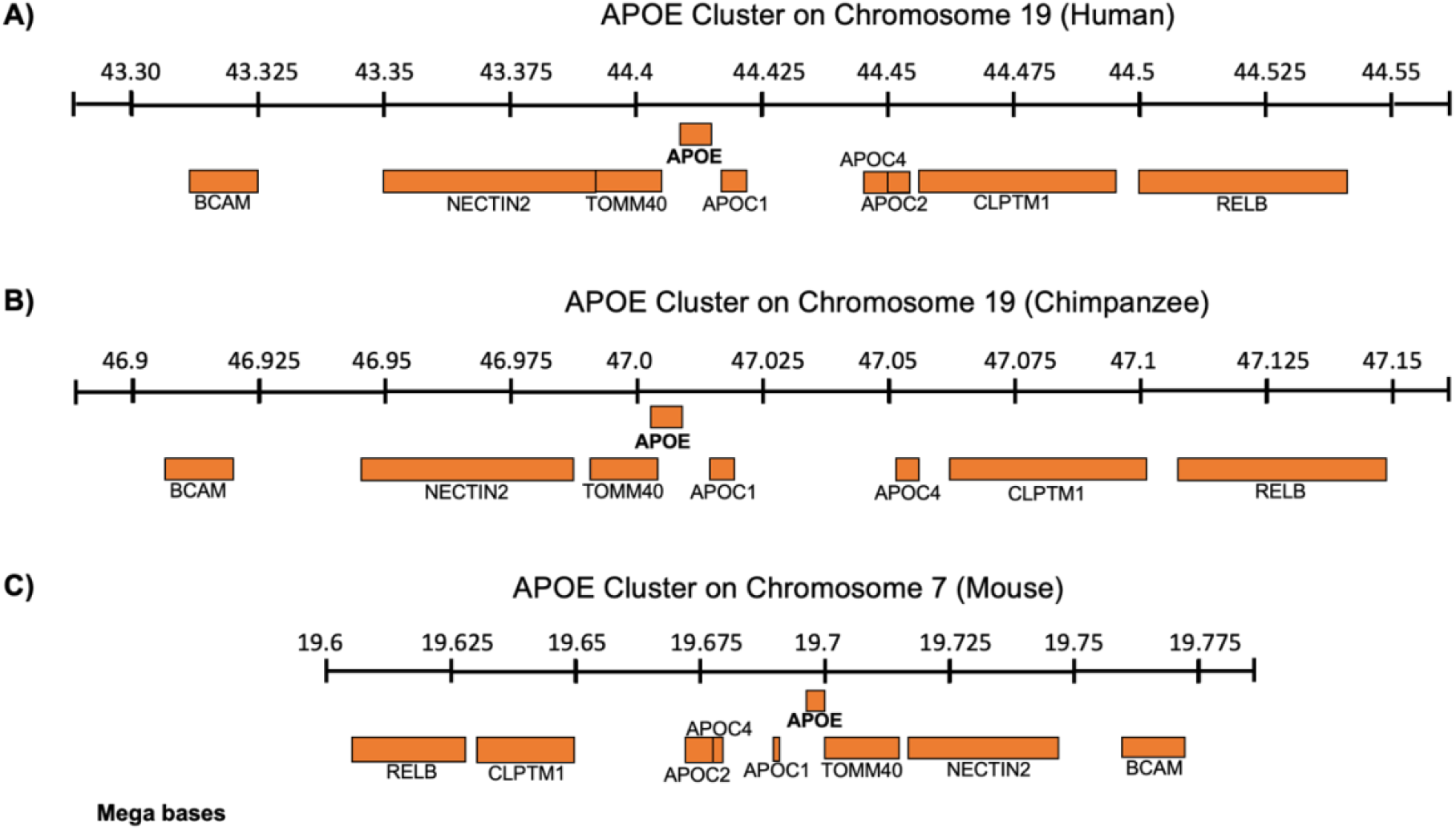
The ApoE gene cluster found of chromosome 19q13.32 for **A)** human and **B)** chimpanzee which lacks ApoC2. The **C)** mouse ApoE gene cluster is located on chromosome 7qA1 with inverted synteny.

Our findings add to a growing body of evidence that ApoE isoforms exert substantial developmental influence on neuronal architecture, well before the onset of neurodegeneration. The observation that chimpanzees, despite having a sequence similar to ApoE4, do not develop AD, may therefore be due not only to protective amino acid substitutions like T61 but also to species-specific differences in gene regulation, lipid metabolism, and astrocyte-neuron signaling. Importantly, the absence of AD-like pathology in aged chimpanzees despite structural similarity to human ApoE4 underscores the limitations of residue-based assumptions about isoform function. Future studies of the chimpanzee ApoE knock-in mouse line across developmental and aging timepoints could reveal when and how these differences emerge.

## Supplemental Figures

**Supplementary Figure 1:**
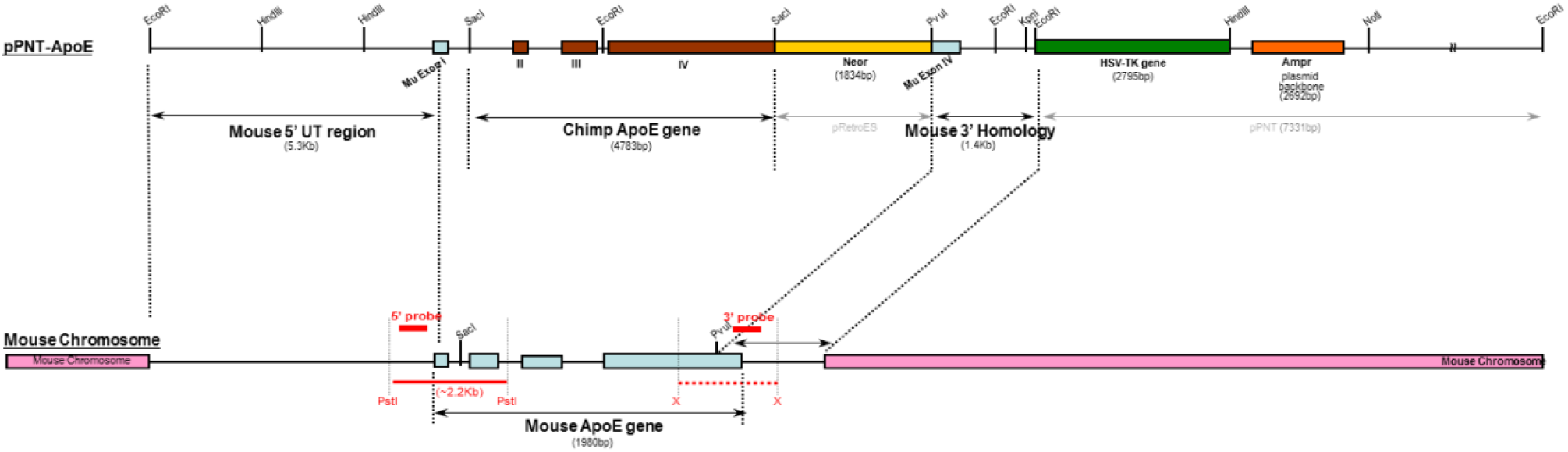
Schematic for targeted replacement of chimpanzee ApoE into C57BL6 mice.

**Supplementary Figure 2:**
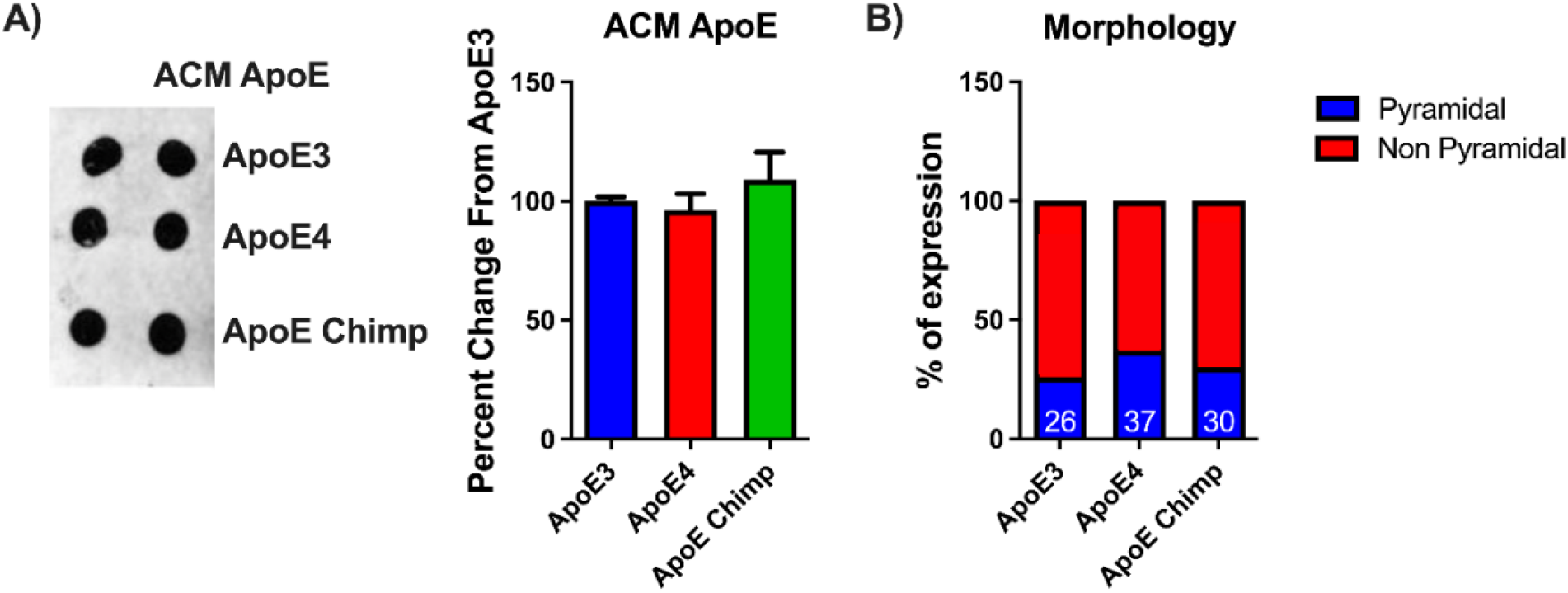
ApoE levels and neuronal subtypes did not differ by ApoE isoform. **A)** Dotblot showing equal levels of ApoE found in ACM and **B)** percent of neuronal subtype expressed in E18 rat neurons in response to ApoE ACM for over 16 individual culture dishes. A) Statistics by one-way ANOVA with Tukey’s posthoc or Kruskal-Wallis. B) Statistics by Chi-square test corrected by Bonferroni.

**Supplementary Figure 3:**
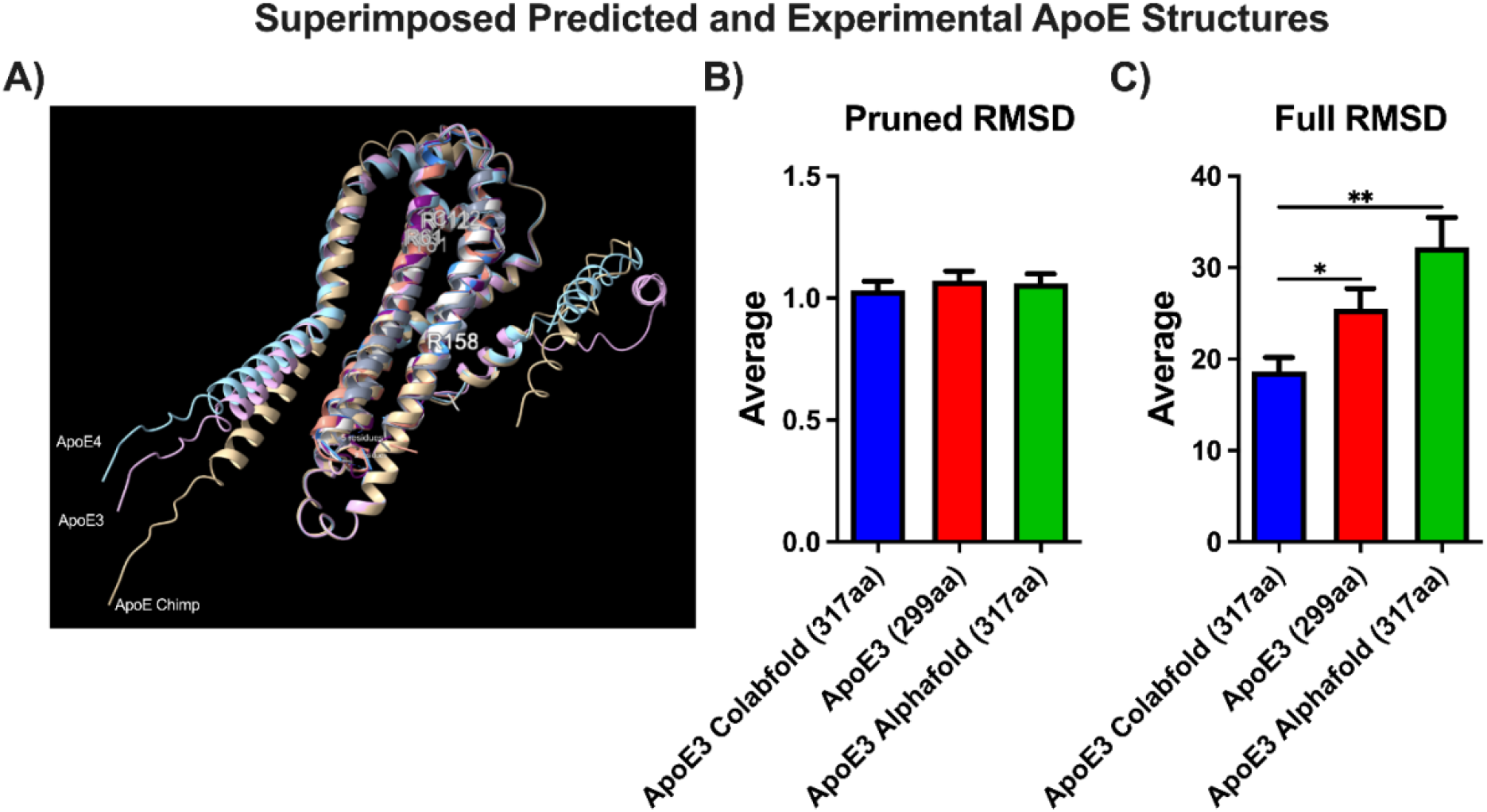
Alignment values generated using matchmaker in ChimeraX in reference to the predicted ApoE3 structure from Colabfold. Statistics by Welch’s ANOVA with Games-Howell posthoc test. *p<0.05, **p<0.01.

**Supplementary Figure 4:**
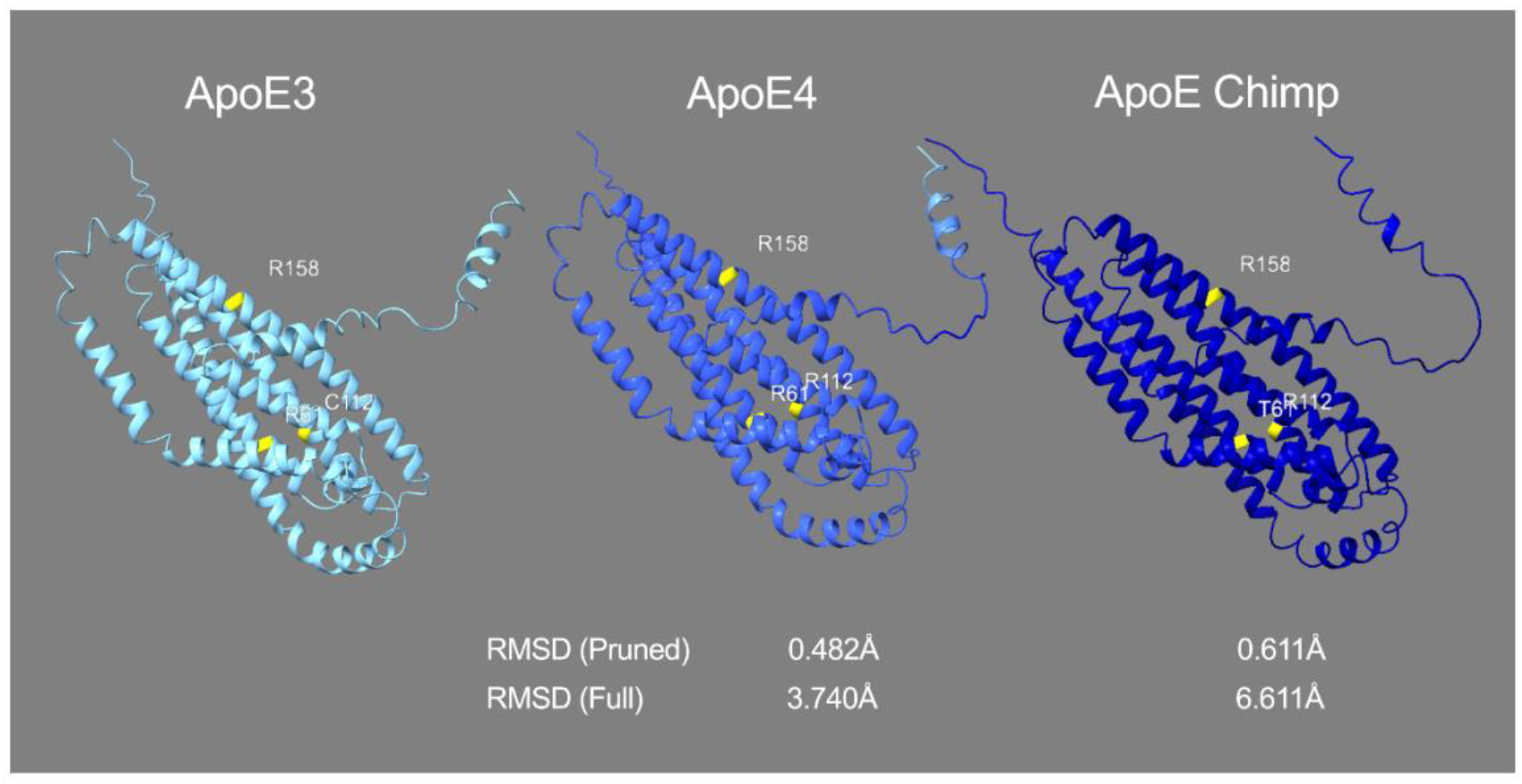
Comparison of precursor ApoE3, ApoE4, and ApoE Chimpanzee structures. Key residues at positions 61, 112, and 158 are highlighted in yellow. RMSD values (pruned and full) were calculated relative to ApoE3. Structures are colored by full RMSD gradient relative to ApoE3 to indicate structural deviation; darker indicates higher shifts in atom positions.

**Supplementary Figure 5:**
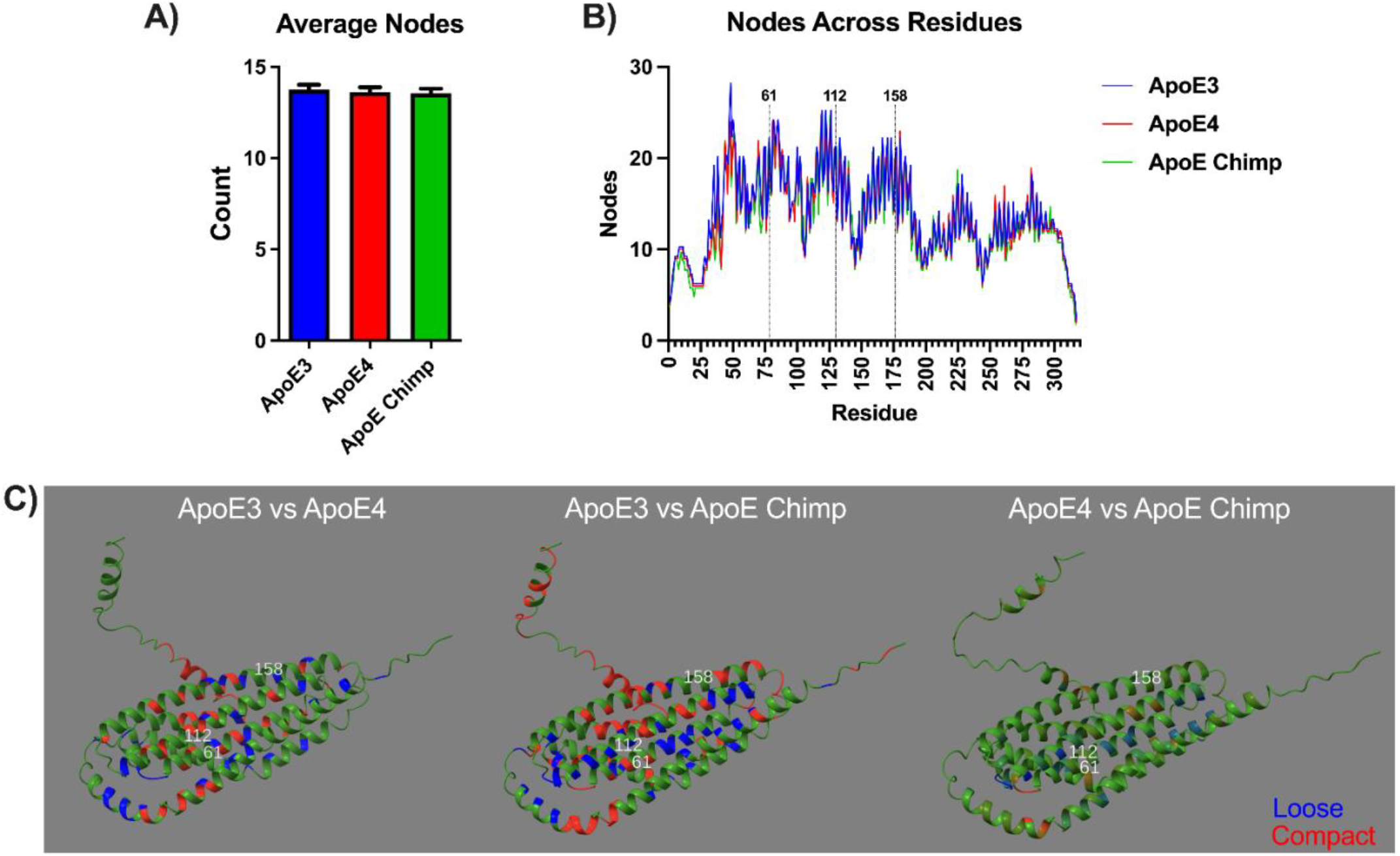
Amino acid connectivity for precursor ApoE3, ApoE4, and chimpanzee ApoE based on residue-level graph connectivity. **A)** The average number of nodes per ApoE isoform using a 10Å cutoff. **B)** The number of nodes at each amino acid for precursor ApoE. Annotations (61, 112, and 158) are based on the mature 299 amino acid sequence. **C)** Reference structures of ApoE3 or ApoE4 in green colored by the Δnode degree for ApoE4 or ApoE chimpanzee. Red indicates increased node connectivity while blue represents decreased node connectivity.

## Acknowledgements

We thank Patrick Sullivan at Duke University for generous donation of immortalized ApoE3 and ApoE4 astrocyte lines. We thank Richard Sprott for supporting the generation of the targeted replacement chimpanzee mouse. BioRender was used for the generation of figure 1.

## Ethics Declarations

All of the authors declare no competing interests.

## Data Availability

The authors declare that the data supporting the findings of this study are available within the paper and its Supplementary Information files. Raw data files may be requested in other format from corresponding authors.

## Author Contributions

Max A. Thorwald (Conceptualization, Data curation; Formal Analysis; Investigation; Methodology; Validation; Visualization; Writing – original draft; Writing – review & editing); Mafalda Cacciottolo (Conceptualization, Data curation; Formal Analysis; Investigation; Methodology; Validation; Visualization; Writing – original draft; Writing – review & editing); Todd Morgan (Conceptualization; Funding Acquisition; Resources; Supervision; Writing – review & editing); Caleb E. Finch (Conceptualization; Funding Acquisition; Project Administration; Resources; Supervision; Writing – original draft; Writing – review & editing).

